# OmeSim: a genetics-based nonlinear simulator for in-between-ome and phenotype

**DOI:** 10.1101/2024.03.10.584320

**Authors:** Zhou Long, Qingrun Zhang

## Abstract

**Motivation:** Deciphering genetic basis of complex traits via genotype-phenotype association studies is a long-standing theme in genetics. The availability of molecular omics data (such as transcriptome) has enabled researchers to utilize “in-between-omes” in association studies, for instance transcriptome-wide association study. Although many statistical tests and machine learning models integrating omics in genetic mapping are emerging, there is no standard way to simulate phenotype by genotype with the role of in-between-omes incorporated. Moreover, the involvement of in-between-omes usually bring substantial nonlinear architecture (e.g., co-expression network), that may be non-trivial to simulate. As such, rigorous power estimations, a critical step to test novel models, may not be conducted fairly.

**Results:** To address the gap between emerging methods development and the unavailability of adequate simulators, we developed OmeSim, a phenotype simulator incorporating genetics, an in-between-ome (e.g., transcriptome), and their complex relationships including nonlinear architectures. OmeSim outputs detailed causality graphs together with original data, correlations, and associations structures between phenotypic traits and omes terms as comprehensive gold-standard datasets for the verifications of novel tools integrating an in-between-ome in genotype-phenotype association studies. We expect OmeSim to enable rigorous benchmarking for the future multi-omics integrations.

**Availability:** https://github.com/zhoulongcoding/OmeSim

## Introduction

Simulators play a critical role in assessing the performance of newly developed computational tools, especially when comparing a new tool to the state-of-the-art. This is particularly important for tool developers in the field of statistical genetics with the aim to discover the genetic basis of complex traits. Additionally, researchers need genotype-based phenotype simulators to assist study-design in which the power estimation has to be carried out (Wang *et al*., 2014). As such, modern genotype-based phenotype simulators have been developed from time to time (Ji *et al*., 2021).

With the availability of modern biobanks providing comprehensive genetic and phenotypic data (Tryka *et al*., 2014), more ambitious statistical methods have been devised. Accordingly, the corresponding simulators have also been developed. For instance, many recent simulators support the generation of multiple correlated phenotypes (Meyer and Birney, 2018; O’Reilly *et al*., 2012; Porter and O’Reilly, 2017), and some simulators support the epistatic effects between causal genetic factors (Fernandes and Lipka, 2020; Reidenbach *et al*., 2021).

Along with the advancement of high-throughput data generation towards multi-scale -omics, researchers now have access to various -omics data including transcriptomes (Ardlie *et al*., 2015), methylomes (Stunnenberg *et al*., 2016), proteomes (Edwards *et al*., 2015) etc., referred to as in-between-omes hereafter. These data exposed the challenges of handling correlations between in-between-ome terms (e.g., co-expressions in the transcriptome), however they also provide opportunities (Wainberg *et al*., 2019) These data have triggered development of sophisticated tools leveraging in-between-omes to characterize the genetic basis of complex traits. Such tools usually adapt complicated machine learning models that may consider nonlinear analysis as well as causality assumptions or inference (Muneeb *et al*., 2022; Meijering and Gianola, 1985; Sailer and Harms, 2017; Bao *et al*., 2020; Basu *et al*., 2018; Lee *et al*., 2016). However, there are no standard simulators to benchmark the performance of the newly developed tools, leaving authors to develop different *ad hoc* simulations tailoring to their works. Partly due to the complexity of multi-scale -omics data and their tricky dependence structure, the design of simulations may have to be somewhat arbitrary and controversial. Therefore, a standard tool of simulating phenotypes and an in-between -omics data that explicitly exposes the multi-scale genetic architecture including the causal relationship between terms is urgently needed.

Based on our recent experience of conducting *ad hoc* simulations involving nonlinear relationships in genomic and transcriptomic data (He, Li, *et al*., 2023; Kossinna *et al*., 2022; Cao *et al*., 2022), here we developed a comprehensive framework, OmeSim, that simulates phenotype based on genotype and an in-between-ome (e.g., transcriptome) using a broad spectrum of genetic architectures including nonlinear relationship and multiple causality assumptions and correlation structures. OmeSim also outputs detailed gold-standard datasets in the format of both text files and automatically generated graphic representations (for the full datasets and for each trait and pathways).

In the rest of this paper, we describe the design principles, underline statistical models and implementation details in **Methods**. The application related issues, i.e., input/output specifications, examples of downstream utilization, as well sample visualizations are presented in **Results**. Future extensions and several design considerations are elaborated in **Discussion**.

## Methods

### 2.1. Design and implementation principles

OmeSim is designed to generate phenotypic traits and an in-between-ome based on a population of genotypes, integrating complicated relationships between genotype, an in-between-ome, and phenotype. Based on the input datasets, i.e., genotype data, genes models, and pathways, the OmeSim-generated phenotype and in-between-ome are directed by simulated causality graph and detailed quantitative models. Users may specify preferred parameters to control this process (outlined later in this section and fully described in OmeSim’s **Users’ Manual**). Such relationships will be fully recorded as correlations and associations between all participating terms (genetics, an in-between-ome, and traits).

The design of OmeSim is structured into *high-level* architectures and *low-level* models. More specifically, the high-level architectures concern the relationship between genome, phenotype and the in-between-ome, which contain four models including Causality, Pleiotropy, Reverse-causality (He, Antonyan, *et al*., 2023; He, Li, *et al*., 2023; Cao *et al*., 2022; Cao, Ding, *et al*., 2021), and Hybrid. The high-level architecture also specifies the variance components explained by the response variables (e.g., the phenotypic variance component explained by the genetic variants, variance explained by other traits, and level of noise terms). In contrast, the low-level models specify the contribution of each term and the relationship between terms (e.g., linear model based on additive functions and nonlinear models based on epistatic, heterogeneous, compensatory, and hybrid functions (Cao, Kwok, *et al*., 2021; Cao *et al*., 2022)). This design enables separation of concerns so that the users can specify different level of parameters accordingly.

In its implementation, OmeSim first loads necessary data such as files providing genotype, gene models, and pathways (**Figure 1A**). Next OmeSim generates a *causality graph* recording contributing factors of each term (genetic variants, gene expressions and traits) based on user-specified parameters directing high-level architectures (**Figure 1B**). Note that, the trans-genetic variants and gene expressions contributing to a focal gene expression or trait will be collected from the same pathway(s) to promote shared component to reflect the real biological networks (Su *et al*., 2021). Then OmeSim iteratively calculates values of the gene expressions and traits contributed by genetics and other terms using low-level models (**Figure 1C**). Note that iterations are needed as the source terms generating a target term may not be finalized in the initial step. After calculating the values, OmeSim generates the gold-standard datasets (*correlations* between terms and *associations* between genetics and terms) (**Figure 1D**) and then adds infinitesimal and noise terms to finalize the ultimate values of gene expressions and traits. Finally, users can use the function *Causality* to query the actual causality relationship between specific terms of interests using the previously generated causality graph (**Figure 1F**).

**Figure 1:**
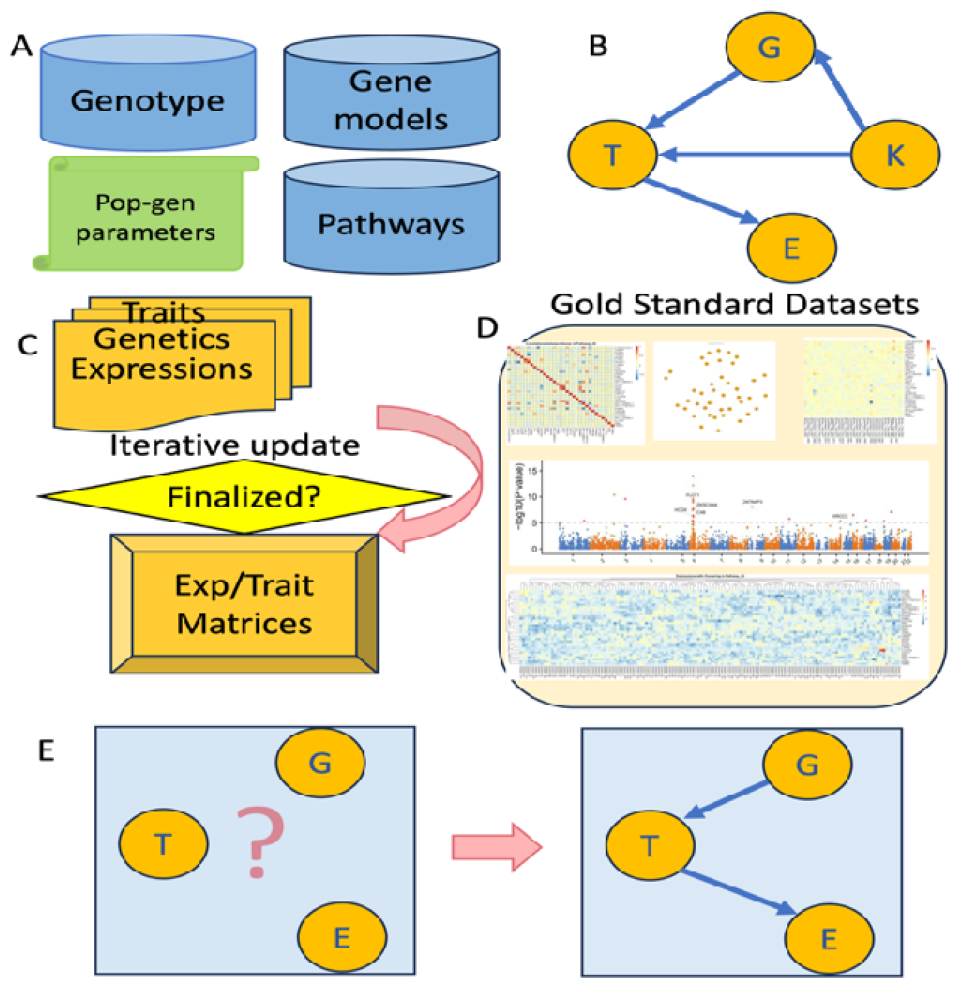
The pipeline of OmeSim. **A**) Input data. **B**) A Causality graph of all terms involved are generated to direct the simulations. **C**) Values of terms are iteratively calculated. **D**) Gold-standard datasets including the in-between-ome/trait values, correlation structure, associations, and causality graphs for each trait and each pathway are output before adding noise and infinitesimal terms. **E**) One may query causality relationship between specific terms.

### 2.2. Notations

Notation-wise, the genotype is denoted as an *N* × *M* matrix ***G***, where *N* is the number of individuals, and *M* is the number of genetic variants. *G*_*i,m*_ denotes genotype of the *i*-th individual at the *m*-th variants. ***G***_*\*,m*_ denotes the vector of genotype of the *m*-th location for all individuals, and ***G***_***i***,*\**_ denotes the vector of genotype of the *i*-th individual for all variants. Similarly, the in-between-ome data (e.g., transcriptome) is denoted as an *N* × *K* matrix ***Z***, where K is the number of terms (e.g., number of genes). Again, *Z*_*i,k*_ denotes the expression value of *i* -th individual on the *k*-th gene. ***Z***_*\**,***k***_ denotes the vector of expressions of the *k*-th gene for all individuals, and ***Z***_***i***,*\**_ denotes the vector of expressions of the *i*-th individual for all genes. The phenotype is denoted by an *N* × *T* matrix ***Y***, where *T* is the number of traits. Again, *Y*_*i,t*_ denotes the phenotypic value of *i*-th individual on the *t*-th trait. ***Y***_*\**,***t***_ denotes the vector of values of the *t*-th trait for all individuals, and ***Y***_***i***,*\**_ denotes the vector of values of the *i*-th individual for all traits.

As we used transcriptomes to demonstrate the models, in the rest of the paper, “transcriptome” will be used to replace “in-between-ome” for simplicity. However, the models apply to any other in-between-ome.

### 2.2. High-level architectures

The high-level architectures specify the causal relationship between contributing terms and the target term, detailed below. Note that in the formulas below, for simplicity, we do *not* distinguish cis- and transgenetic variants, although they are separated in the implementation (**Users’ Manual**). The cis- and trans- variants are denoted by the subindex “*M*”.

#### Causality model

Genetic variations are the cause of transcriptomic variations, which in turn causes phenotypic changes. Specifically, we have ***Z***_***k***_ = *f*_*c*_(***G***_***M***_) + *ϵ*, and ***Y***_***t***_ = *g*_*c*_ (***G***_***M***_, ***Z***_***k***_) + *ϵ*. The subindex “*c*” standards for “causality”.

#### Pleiotropy model

Genetic variations cause the transcriptomic variations as well as phenotypic changes via independent models. Specifically, we have **Z**_**k**_ = *f*_*p*_(***G***_***M***_) + *ϵ*, and ***Y***_***t***_ = *g*_*p*_(***G***_***M***_) + *ϵ*. The subindex “*p*” standards for “pleiotropy”. Please note that, here “independent” means the functions *f*_*p*_(. ) and *g*_*p*_(. ) are independent. Since both ***Z***_***k***_ and ***Y***_***t***_ are dependent on ***G***_***M***_, they are still correlated (and the actual correlation will be quantified by OmeSim during the simulation).

#### Reverse-Causality model

Genetic variations are the cause of phenotypic changes, which in turn causes transcriptomic variations. Specifically, we have *Y*_*t*_ = *g*_*r*_(*G*_*M*_) + E + \epsilon, and *Z*_*k*_ = *f*_*r*_ (*G*_*M*_, *Y*_*t*_) + E. The subindex “r” standards for “reverse-causality”.

#### Hybrid model

The above three models assume that all the terms in phenotype (traits) or transcriptome (genes) follow the same relationship. However, in practice, it is likely that a subset of genes expressions is influenced by the change of traits; and a subset of traits may be altered by other gene expressions. As such, we provided a mixture of the above three models, referred to as “Hybrid”, which is actually the default of OmeSim. This could be randomly generated by OmeSim or specified by the users.

#### Causality graph

The above-simulated high-level causal-relationship between genes and traits are recorded in a causality graph output by OmeSim during the simulation process. Please see **Figure 2A** for a subset of the global picture, **Figure 3A-D** for the example of a trait, and **Figure 4B** for the example of pathway. OmeSim also provides a query function “*Causality*” to check the causality relationship between any specified triples of terms (**Users’ Manual**).

**Figure 2:**
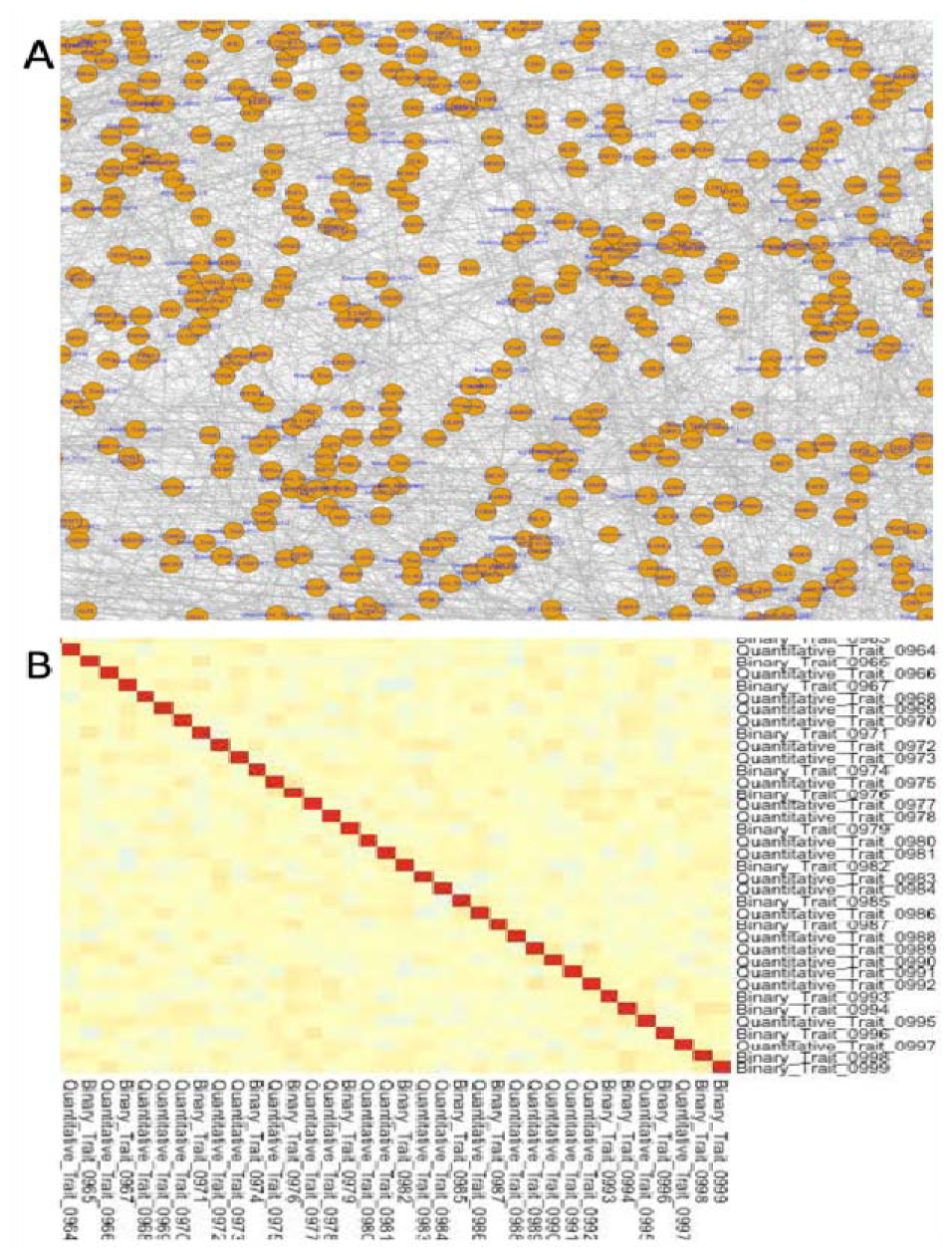
Examples of full views. **A**) A part of the OmeSim generated global causality graph of all traits and expressions. **B**) The bottom-right conner of the trait-by-trait correlation heatmap.

**Figure 3:**
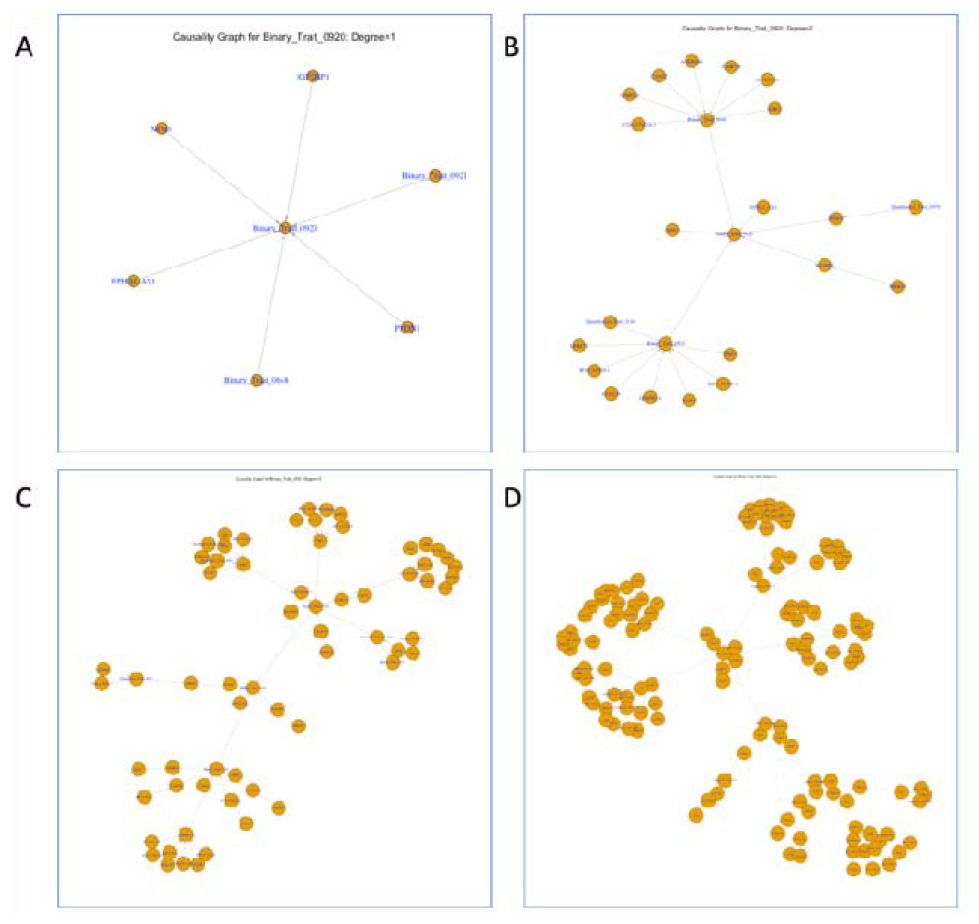
Example sub-causality graphs for a simulated trait (#0920) **A**) Degree = 1. **B**) Degree = 2. **C**) Degree = 3. **D**) Degree = 4.

**Figure 4:**
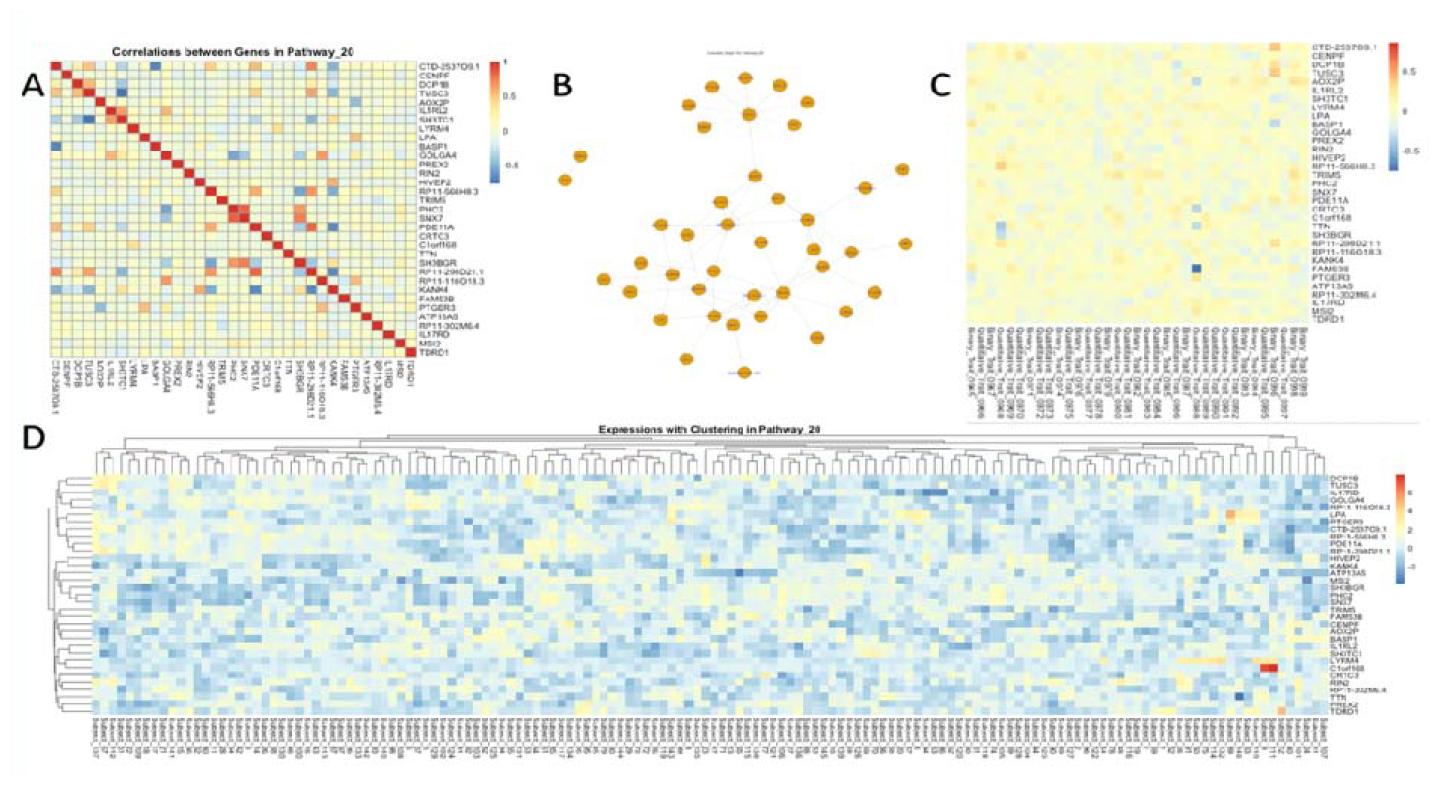
Pathway-specific visualization (an example of simulated Pathway #20). **A**) Heatmap visualizing correlations between all genes within the pathway **B**) Sub-causality graph within the pathway. **C**) Heatmap visualizing correlations between genes within the pathway and simulated traits (only 35 traits in the right-most part of the figure are shown. **D**) The expression data in all subjects with bi-clustering of genes and subjects.

### 2.3. Specific statistical models

The low-level specific models concern the actual definition and implementation of focal terms, *g*_*a*_ (. ) and *f*_*a*_(. ), where “*a*” indicates any subindex *c, p*, or *r*. There are five models, for which users can specify the proportion of them by setting up their distribution parameters (**Users’ Manual)**. The models are described below.

#### Genotype representation

On the finest scale, OmeSim incorporates three coding schemas to encode genotypic alleles, representing three alternative genetic models: *additive, dominant*, and *recessive*. More specifically, OmeSim will use 0, 1, and 2 to represent genotype a/a, a/A and A/A (where “a” and “A” stand for reference and alternate alleles respectively), reflecting well adapted *additive* model. Alternatively, (0, 2, 2) or (0, 0, 2) may be used to adapt *dominant* model (heterozygous mutant = homozygous mutant) and *recessive* model (heterozygous mutant = reference). These different coding indeed will bring significant difference in the analysis of genetic components of phenotypic traits (Huang and Mackay, 2016). OmeSim accepts numerically coded variants in CSV file or raw genotype (i.e., coded by A, C, G, T) together with specified parameter for the selection of genetic models (i.e., *additive, dominant*, and *recessive*) for OmeSim to convert to corresponding representations (**Users’ Manual**).

#### Linear and interactive models

OmeSim provides an extensive flexibility of defining how multiple terms impact the outcome (gene expression or phenotypic traits) jointly. First, *Linear models* are the simplest ones to form the relationship: *f*(***T***_***M***_) = *Σ β* _*i*_ *T*_*i*_, where T represent the matrix of terms composed of genetics (*G*), expression (*Z*), and/or traits (*Y*). Three nonlinear models are defined as *epistatic, compensatory*, and *heterogenous*. Intuitively, in epistatic models, one of the terms acts as the “switch” that enables other terms to be effective; in compensatory models, one term can compensate the effect of the other; in heterogenous models, any of the terms’ change can independently cause the changes of the target term and multiple terms’ change will not lead to higher result. When there are only two terms and the terms are genotypes or binary traits (represented by “true” or “false”), the above three models are equivalent to the standard logical gates AND, XOR (= exclusive OR) and OR. When the terms are not binary traits, i.e., genotypes coded in a schema mentioned above, quantitative gene expressions or traits, OmeSim use the following process to ensure that they are not simply converted into binary (that may cause the future calculations relying on them not meaningful). OmeSim first re-scales the vector’s mean to be 0.0 and variance to 1.0, which is a common practice in genetic analysis (Yang *et al*., 2011). Then the quantitative measurement will be considered “false” if lower than zero or “true” if higher than zero. Then OmeSim defines operations similar to standard logical gates to ensure that the outcome will be a negative value if “false” or a positive value if “true”. This operation ensures that the resulting vectors remain quantitative until the final conversation to binary (if it is a binary trait). The detailed definitions are given below:

#### Epistatic model

(which is similar to the AND logical operator) *f*(*T*_*M*_) = *Λ T*_*i*_ (i = 1,2, …, m) where the operator *T*_1_ Λ *T*_2_ is defined as *T*_2_ if both *T*_1_ and *T*_2_ are positive; or min (*T*_1_, *T*_2_) if both are negative.

#### Compensatory model

(which is similar to the XOR logical operator, and is defined on two terms only): *T*_1_ *⊕ T*_2_ is defined as: (1) if they have the same sign, the result is max (*T*_1_, *T*_2_) if both are positive; or -min (*T*_1_, *T*_2_) if both are negative. (2) if they have different sign, then the negative one, which must be the smaller one, i.e., min (*T*_1_, *T*_2_) is the result.

#### Heterogenous model

(which is similar to the OR logical operator): *f*(*T*_*M*_) = V *T*_*i*_ (*i* = 1,2, …, *m*) where the operator *T*_1_ V *T*_2_ is defined as *T*_2_ if both *T*_1_ and *T*_2_ are negative; or max (*T*_1_, *T*_2_) if both are positive.

#### Compound models

More complex models that are compounds of the above four models (i.e., *linear, epistatic, compensatory*, and *heterogenous*) can also be specified. This is achieved by recursive use of these models. For instance, one can first define a “super-gene” depending on three genotypic variants using *additive* model, and then consider this “super-gene” as one term in another function, say *epistatic*. If the users prefer to use *compound* model, however, do not provide specific instructions on how the detailed steps are defined (which is common as one may not manually define all operations), OmeSim will generate such compound protocol randomly based on an internal rule (details presented in **Users’ Manual**.)

#### Weights of different contributors

As stated, in the above functions, the terms ***T***_***M***_ represent the matrix of terms composed of genetics (*G*), expression (*Z*), and/or traits (*Y*). The users can specify their relative importance by providing different weights. For gene expressions, cis- and trans-regulatory variants enjoy two separate weights. For instance, regional heritability (Grundberg *et al*., 2012) can be reflected by the contribution of cis-genetic weights.

#### Noise and infinitesimal terms

In the above models, the variance components explained by the actual biological input are modeled by the focal terms (*g*_*a*_(. ) and *f*_*a*_(. ), where “*a*” indicate any subindex *c, p*, or *r*, and the residual *ϵ* is randomly generated (directed by the parameters specified by the users or the defaults). Here the residual term ϵ may be entirely random (based on user-specified or default variance components), which could be called “*noise*” term, or may be structured to the sum of a truly random term and a genetic background term. The genetic background term may also be called as an *infinitesimal* (Cui *et al*., 2023) term, the contribution from genetic background containing many small genetic effects. This aggregated term may also be called “polygenic” (MATHER, 1943) term. OmeSim implements the addition of such an infinitesimal (or polygenic) term by generating an additive term aggregating many genetic variants’ tiny contributions. In particular, OmeSim samples a large number of genetic variants (default = 5,000, adjustable by the user) and assigns a coefficient (randomly drawn from standard normal distribution N(0,1)) to each genetic variant. The sum of these variants’ effects, re-scaled by the variance component specified by the users (or randomly generated by OmeSim based on user-specified or default ranges) will be added to the target term.

#### Converting quantitative values to binary traits

The above models, including functions *g*_*a*_ (. ) and *f*_*a*_(. ), as well as the noise term and potentially a polygenic term, simulate gene expressions phenotypic traits are naturally quantitative. When it is a trait and the users prefer to set it to a binary trait, reflecting the “case” and “control” status in disease studies, OmeSim will convert them into binary traits. This is done through two alternative methods: The first is the standard logistic regression model (Ott, 2023), in which an odd will be calculated by 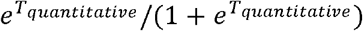 ), equivalent to the logistic model specifying 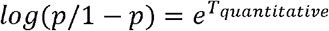 (where 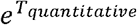 is the “raw” quantitative outcome). Then if the odd is higher than a cutoff (specified by the user, or is 0.5 by OmeSim default), the binary trait will be set up to 1, otherwise 0. The second method is to use a simple liability model (Zuk *et al*., 2012), i.e., the binary trait will be set up to 1 or 0 depending on whether the raw quantitative outcome is higher than a prespecified cutoff (specified by the user or is the median by OmeSim default). The selection between the above two methods will be randomly assigned by OmeSim by the user specified probability distributions (Users’ Manual). In the event the user prefers one method against the other, simply setting the probability of that method to be 1.0 (and the other 0.0) will achieve the goal. Additionally, how many traits are simulated to be quantitative or binary will also be specified by the user in a similar way.

#### Correlations and associations between terms

To assist *type-I error* (or any other false positive analysis) and *power* analysis when benchmarking a newly developed tool, OmeSim outputs “gold-standard” measurement of simulated terms. These include five actual correlation matrices between the terms. They are correlations between gene expressions ( *K* × *K* ), correlations between traits ( *T* × *T* ), correlations between traits and expressions ( *T* × *K* ), correlations between gene expressions and their contributing genetic variants (*K* × *M*), and correlations between genetic variants and traits and their contributing genetic variants (*T* × *M*). These are called “gold-standard” because that they are calculated using the calculated values **<u>before</u>** adding the residuals terms including both noise and infinitesimal (if any). Without a simulator, the analytic tools can only calculate such correction matrices using real data (e.g., a co-expression network) that contains substantial noise. That is why OmeSim outputs the correlation matrices before adding these noise and infinitesimal terms. Additionally, the correlations because of artifact factors such as linkage disequilibrium (LD) (Wang *et al*., 2023; Slatkin, 2008) is distinguished from actual biological correlations as non-contributing genetic variants will not be assessed correlations to expressions or traits. These gold-standard files, together with the causality graph generated from the high-level architectures, will lay a solid ground for the assessment of the statistical performance of a newly developed tool.

#### 2.4. Other implementation considerations

OmeSim focuses on the relationship of traits and an in-between-ome, therefore does not simulate genotypes. Users may use standard reference panels, such as the 1000 Genomes Project or the genotype file in a GWAS dataset as the input genotype file. If some population genetic properties are desired, one may use other simulators such as MS (Ewing and Hermisson, 2010), SLiM (Haller and Messer, 2019) or Hapgen (Su *et al*., 2011), by which specific evolutionary scenario or population trajectory are supported.

#### Selection of genetic variants

Given the full genotype file, OmeSim can simulate expressions and phenotypic traits using the user specified genetic variants. In the event the user does not specify concrete variants in the form of specific locations, OmeSim will select variants based on the user’s instructions, such as the number of variants and their minor allele frequency (MAF). Linkage disequilibrium (LD) between variants may also be considered by specifying the minimal distance between adjacent variants. The detailed specifications are presented in **Users’ Manual**. *Shared pathways and correlations between traits/expressions*. OmeSim-simulated expressions and traits are correlated because of the following two reasons: first, the expressions contributing to traits, as well as common genetic components directly generate such correlations. Second, to mimic the internal structure caused by pathways, all trans-genetic variants and gene expressions contributing to a trait or an expression will be selected from the same pathway(s). This design will lead to the “gene-modules” (Langfelder and Horvath, 2008) frequently observed in co-expression analysis.

#### Visualization

To facilitate both a global view and detailed inspections, visualizations are provided in various resolutions. Heatmaps are provided to view the overall correlations between all terms. The overall full causality graph and the pathway-specific and trait-specific sub-causality graphs are also be provided. Based on the user’s queries, detailed causal relationship and genetic associations will also be generated. Please see **Users’ Manual** for detailed instructions and **Section 3.2 An example use-case** for illustrating instances.

## Results

### The input and output files

OmeSim requires three data input files: a genotype file, a gene-model file to specify the locations of genes, and pathways file specifying which genes are in the same pathway. Importantly, all parameters are provided in a configuration file, specifying various user-specified requirements OmeSim requires the input genotype file in CSV format and will support VCF and PLINK soon. The gene model and pathways are formatted in CSV files. The configuration file has to follow our specified format, specifying the required high-level architectures and low-level models described in **Methods**. The full details are described in the **Users’ Manual**.

The output text files are gene expressions and phenotypic traits in a CSV format for specific downstream analysis to test newly developed tools such as TWAS tools (Gamazon *et al*., 2015; Cao, Kwok, *et al*., 2021; He, Li, *et al*., 2023; Cao *et al*., 2022), or co-expression analysis tools (Kossinna *et al*., 2022). Importantly, “gold-standard” files recording causal genetic variants and correlations between terms will also be stored in the output folder. These correlation files, together with the causality graph, play critical roles in such assessments.

### An example use-case

We use the following representative use-case to describe a step-by-step execution of OmeSim.

Suppose one wants to simulate a case-control dataset together with 1,000 (possibly correlated) phenotypic traits and transcriptome for around 7,000 genes based a combination of linear and nonlinear genetic architecture as well as a mixture of all causality models. The starting point will be setting up the configuration file.

The paths to input data files described in the previous section are mandatory parameters specified in this configuration file. The rest parameters are roughly outlined below. The user’s requirements such as the causal variants may be common variants (i.e., MAF > 0.05) and are not in strong LD are also specified in the configuration file. Additionally, the user may prefer the distribution between five detailed models (*additive, epistatic, compensatory, heterogenous* and *compound*) to follow prespecifies probabilities. The user may want to specify that all gene expressions have cis-regulatory effects and all traits contain genetic effects as well as the probability of their contributors including other gene expressions or traits, and infinitesimal term as well. The user may also prefer the relative weights between genetic, expression and trait contributions to be specified using distributions. Moreover, the users may specify the number of genetic variants and terms in the models. Besides, many statistical genetic model parameters need to be set up. All these are supplied to OmeSim by the configuration file. Although there are many parameters that the users may or may not use, we provide a default configuration file that is available on the OmeSim GitHub together with the detailed explanations of the meaning of the parameters (**Users’ Manual**).

After running OmeSim, all output text files are in the user-specified output folder. Importantly, OmeSim provides visualization of both global-level and fine-scale data. For global-level structures, OmeSim provides six figures in the root of output folder, displaying global causality graph (part is shown in **Figure 2A**), trait-by-trait correlation heatmap (part is shown in **Figure 2B**), expression-by-expression heatmap, expression-by-trait heatmap, and the original data of expression and traits with clustering, also visualized by heatmaps.

To facilitate investigation of the causal factors any trait, OmeSim also generates sub-causality graphs for each trait. We call all the terms directly causal to the trait “first-degree” and the terms in-directly causal second-, third-, …, degree (as defined in standard graph theory). The first four-degrees sub-causality graphs for a trait is presented in **Figure 3**. To facilitate investigation of the causal factors any trait, OmeSim also generates sub-causality graphs for each trait. We call all the terms directly causal to the trait “first-degree” and the terms in-directly causal second-, third-, …, degree (as defined in standard graph theory). The first four-degrees sub-causality graphs for a trait is presented in **Figure 3**.

To understand the gene-gene and gene-trait interactions related to a particular pathway, OmeSim provides detailed text files and visualizations within each pathway. These include correlations between each pairs of genes of this pathway (**Figure 4A**), sub-causality graph of the genes and their causal traits (if any) (**Figure 4B**), correlations between each gene and all traits (part is shown in **Figure 4C**), and the original data of the expressions in the pathway with bi-clustering (**Figure 4D**). Additionally, causality relationship between individual triples of terms can be generated by OmeSim by an on-the-fly queries. For instance, by asking OmeSim the casualty relationship of the any triples, say [A, B, C], OmeSim can provide their causal relationship one of the three models Causality, Pleiotropy, and Reverse-causality.

The input/output files of the above example are available in the OmeSim GitHub.

## Discussion

In this work, we designed and implemented OmeSim, a genetics-based simulator to generate both an in-between-ome and phenotypic traits. The in-between-ome and the traits are designed to be involved in nonlinear and complex dependency and correlation structures. OmeSim provides user-specified parameters to direct the process of data generation and outputs detailed internal causality and correlation files as “gold-standard” for the benchmarking of novel tools analyzing genetics and omics data, in particular the efforts of discovering genetic basis of phenotypic trait(s) assisted by an in-between-ome.

In the process of simulating the high-level causal architectures, OmeSim does not prevent the generation of cycles. For instance, a gene expression A may contribute to the generation of a trait B, which may impact the expression of A back through some sophisticated cycle. This type of cyclic dependency may cause a problem in data generations. However, we do not present these problems as in real biological systems, such cycles are observed frequently (Tisch *et al*., 2014; Schuhwerk and Brabletz, 2023). So, we leave the system to iteratively update the values along the cycle, mimicking the natural process in organisms. Computationally, by suggesting users to specify that all gene expressions and traits have non-zero genetic contributions, OmeSim ensures that all terms will be initialized at least in the second round of calculation (as the first round will add the genetic contributions to all terms). A question is that in the computations how many that cyclic dependency may occur and whether the values of terms in a cycle indeed converge. We have tested this issue in our example data and found that 93% of genes are not in a circle and converged after 10 rounds of iteration.

There are alternative ways to calculate infinitesimal terms using a GBLUP-like model (Gao *et al*., 2012). For instance, if the users specify an *N* × *N* relationship matrix, *R*, between all participating individuals (e.g., a genomic relationship matrix, or GRM (Yang *et al*., 2011), that is frequently used in GWAS). Using the matrix, one can calculate the polygenic terms ***U***= ***B****r*, where r is a vector sampled as i.i.d from standard normal distribution *N*(0,1), and *B* satisfies ***R***= ***BB***^***T***^. Note that, if the relationship ***R*** is calculated by the standard way of calculating the co-variance, i.e., (***GG***^***T***^)/***M***, then the above method is mathematically equivalent to the current simple method implemented in OmeSim. However, this way allows the flexibility for the users to calculate the relationship matrix differently. Nevertheless, OmeSim used a simpler way for its computational efficiency. If the number of individuals is too large so that large memory is needed to load the relationship matrix, the GBLUP-like method will cause substantial requirement of the computational infrastructure.

Similar to existing simulators, the input genotype file for OmeSim is based on subject-level genotype data, instead of summary statistics, which is becoming prevalent and popular nowadays (Bulik-Sullivan *et al*., 2015). If the goal is to test a novel tool based on summary statistics, one can generate summary statistics (such as Z-scores) using OmeSim simulated traits and the original genotype. Then the whole pipeline of summary statistics-based analysis can be applied.

## Supporting information

Users' Manual

## Author contributions

Z.L. implemented the software, conducted the analysis, and prepared the GitHub and Users’ Manual. Q.Z. conceived the study, supervised the work, and wrote the manuscript with input from Z.L.

