## Supplementary material for "OmeSim: a genetics-based nonlinear simulator for in-between-ome and phenotype": Users' Manual

### User's Manual for OmeSim 1.0

#### 1. Introduction

OmeSim is a comprehensive simulator that simulates molecular omics data (i.e., whole-transcriptome data in its first release) and multiple (possibly correlated) phenotypic traits simultaneously. The immediate application of OmeSim is to support the assessment of novel statistical models and computational tools that aim to integrate multi-omics in the discovery of genetic basis of complex traits.

Unlike most other simulators, OmeSim emphasizes the following features that are becoming the focus of many statistical and computational tools:

- Simulates the whole-transcriptome and the whole-phenome together based on user-specified genotype file, gene file, pathway file, and many parameters. It supports nonlinear genetic models including epistasis, compensatory, heterogenous and the compound combination of them. This is in addition to the standard additive model and the infinitesimal model.
- Supports various causality models (causality, pleiotropy, and reverse causality) at the level of individual terms (i.e., genetic variants, gene expressions, and traits), forming a trait-specific and pathway-specific causality graph (e.g., **Figure 3** in Section Output Files)
- Outputs expression-expression, expression-trait, and trait-trait correlations as well as genetics-expression and genetics-trait associations BEFORE adding noise (= random residual) and in infinitesimal (= contribution from the genetic background) terms.

The genuine correlation and associations, together with the causality graph, serve as the "gold-standard" to assess statistical properties of tools aimed at discovering genetic basis in the presence of complicated linear and nonlinear relationship.

In addition to the function `Simulate` that simulates data, another function `Causality` provide functions to check causality relationship between any three terms under investigation based on the simulated causality graph.

#### 2. A quick start

After downloading the tar ball OmeSim.tar.gz, please just decompress the file by

```
> tar -xvzf OmeSim.tar.gz
```

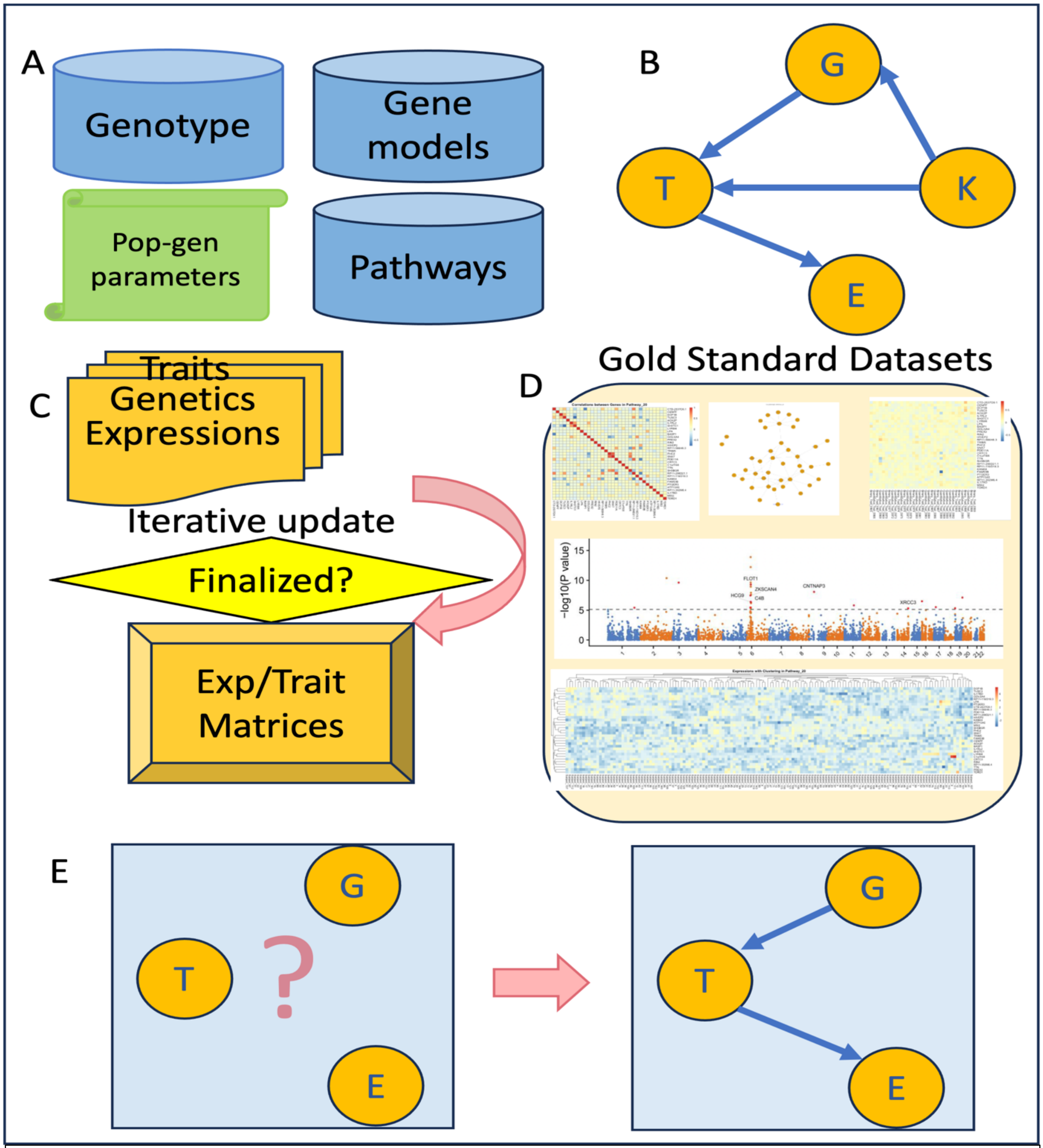


**Figure 1: The pipeline of OmeSim**. **A**) Input data. **B**) A Causality graph of all terms involved are generated to direct the simulations. **C**) Values of terms are iteratively calculated. **D**) Gold-standard datasets including the in-between-ome/trait values, correlation structure, associations, and causality graphs for each trait and each pathway are output before adding noise and infinitesimal terms. **E**) One may query causality relationship between specific terms.

One will see the executable OmeSim.jar, a parameter file (parameter.txt), sample input files, and sample output files. Please modify the paths of the sample input files and the output file folder corresponding to the project folder in your local computer, and type the following commend:

```
> java -Xmx4g -jar OmeSim.jar Simulate -input parameter.txt
```

Then the program will output the sample outcome files in the specified folder. By comparing them to the example outcome files, one can verify whether the program is running smoothly.

#### 3. Design and implementation

OmeSim function `Simulate` adapts a bottom-up strategy to simulate omics (i.e., gene expression in the release 1.0) and phenotypic traits. Depicted in the flow-chart below, OmeSim first loads necessary (genotype, gene models, and pathway) data (**Figure 1A**), then generate a causality graph recording quantified contributing factors of each terms (gene expressions and traits) based on user-specified parameters (**Figure 1B**), and then iteratively calculates values of the gene expressions and traits (**Figure 1C**) contributed by genetics and other terms. After calculating the values, OmeSim will generate the gold-standard datasets (correlations between terms and associations between genetics and terms) (**Figure 1D**) and add infinitesimal and noise terms to finalize the ultimate values of gene expressions and traits (**Figure 1E**). The function `Causality` simply checks the causality relationship using the previously generated causality graph (**Figure 1F**).

#### 4. Input files

For the `Simulate` function: There are four input files: genotype file, gene coordinates file, pathway file, and a parameter file specifying various values of parameters.

- The genotype file should be provided using CSV, PLINK, or VCF formats, specifying genome-wide genetic variants, which are the genetic basis of all omics and traits. It is recommended to use CSV file that is the most simple and efficient format to store genotype information:
  - `Chr,Loc,Genotype_1,Genotype_2,Genotype_3,…, Genotype_n`
  - The genotype above has to be coded using `0` (homozygote reference allele), `1` (heterozygote) and `2` (homozygote alternative allele).
- The gene coordinates file specifies the gene names, the chromosome and start/end locations of all genes. The columns should be separated using commers:
  - `Gene_ID,Chr,Start,End`
- The pathway file should specify the membership of all pathways. The pathway ID and the list of genes are separated by a tab ("\t") and the genes are separated by commers:
  - `Pathway_ID"\t"Gene_1,Gene_2,…,Gene_n`
- The parameter file will be explained in Section "Parameters and their explanations"

Examples of these files are in the download tar ball.

For the `Causality` function, the input are the lines of term triples subjected to the causality check. Each line is composed of three terms separated by commers.

#### 5. Output files

There are three categories of output files for the function `Simulate`:

First, simulated expression and traits will be recorded as `expression.csv` and `traits.csv` in the output folder. In these two files, each column represents the values (of expressions or traits) for an individual; and each row represents the values of all individuals for an expression or a trait. Together with the input genotype data, hey are the main data for running other tools aiming to identify the genetic basis of complex traits by integrating expressions.

Second, causality graph will be recorded in `causality_graph.csv` in the output folder. This file serves as the overall picture of the causality relationship. In this file, each row is composed of terms (genes or traits) separated by commers, with the first gene the term being contributed and the rest contributing. If there is only one term, it means that this term has no other contributor, i.e., only contributed by its own cis genetics.

Third, three correlation files, i.e., `corr_exp_by_exp.csv`, `corr_trait_by_trait.csv`, and `corr_trait_by_exp.csv` recorded the gold standard correlations between these terms before adding noise and infinitesimal terms. Additionally, two association files, i.e., `asso_trait_by_genotype.csv` and `asso_exp_by_genotype.csv` recorded the gold-standard associations between terms and genotypes.

Fourth, the heatmap of correlation files and association files, the visualization of the causality graph is also provided in the output folder. The examples are in **Figure 2 and Figure 3**.

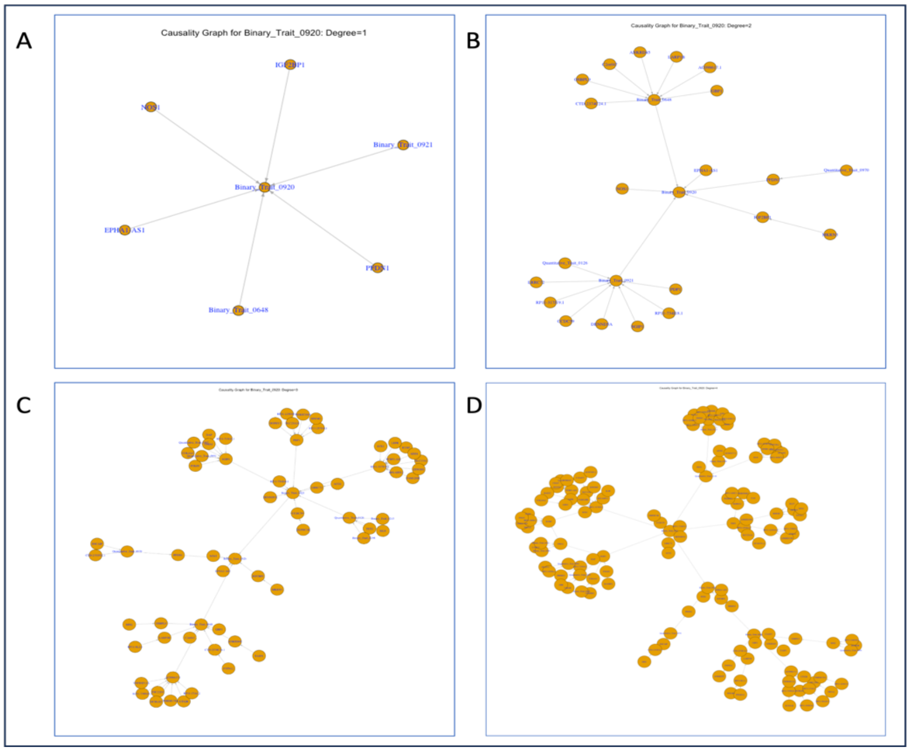


**Figure 2: Example sub-causality graphs for a simulated trait (#0920) A**) Degree = 1. **B**) Degree = 2. **C**) Degree = 3. **D**) Degree = 4.

The output files for function `Causality` are the file of causality models answering the queries lines in the input file together with visualization files (if elected by the input parameters).

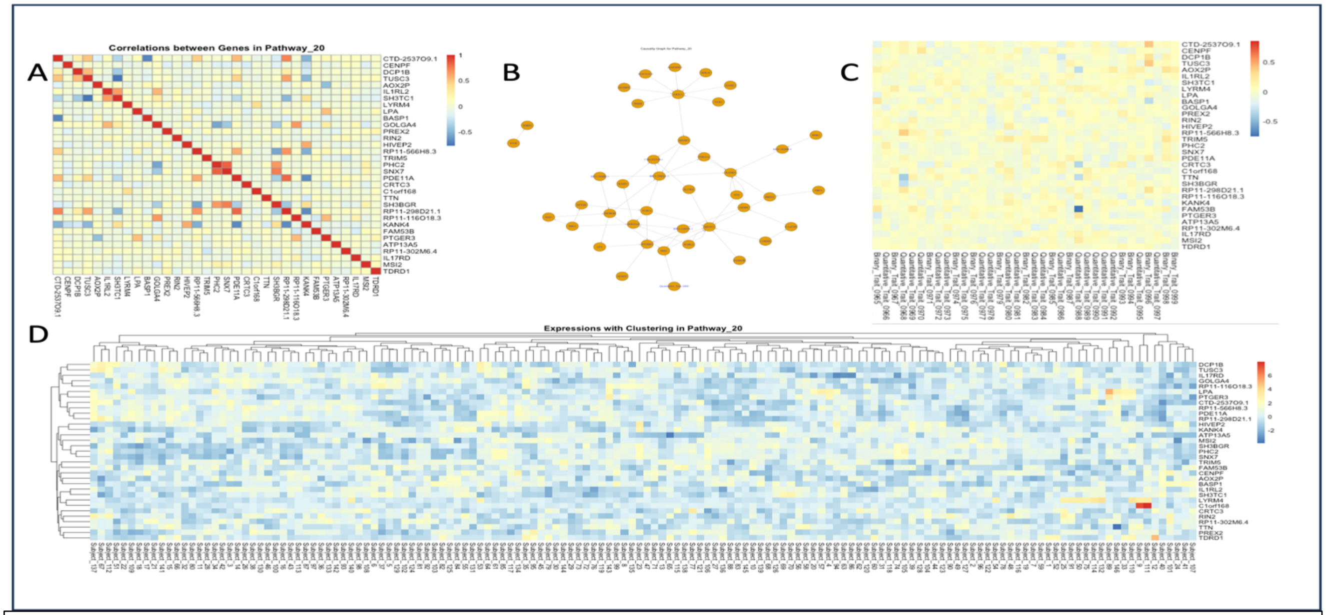


**Figure 3: Pathway-specific visualization (an example of simulated Pathway #20). A**) Heatmap visualizing correlations between all genes within the pathway **B**) Sub-causality graph within the pathway. **C**) Heatmap visualizing correlations between genes within the pathway and simulated traits (only 35 traits in the right-most part of the figure are shown. **D**) The expression data in all subjects with bi-clustering of genes and subjects.

### 6. Parameters

**Parameters for the function `Simulate` can be divided into three categories:**

**Parameters specifying input/intermediate/output files:**

Genotype_File. Full (absolute) path to a file containing genotype information.

Genotype_File_Format. Format of the genotype file. Three formats are supported: "CSV", "PLINK", and "VCF".

Output_Folder. Full (absolute) path to the folder for output files.

Arch_Detailed_File. The intermediate file recording the contributors of each term and their weights. This file will be generated by OmeSim during the computation. Each line specifies the contributors to a term, containing 10 columns:

- ID of the term being contributed (a gene or trait).
- Number of cis regulatory variants.
- Number of trans regulatory variants.
- IDs of genes that the trans regulatory variants located.
- ID of genes whose expressions are relevant.
- ID of traits contributing to the term.
- Weights of the above four categories (cis, trans, expression, and traits).

- `Genetic model (one out of the five supported models, i.e., additive, epistasis, heterogenous, compensatory, and compound).`
- `The proportion of infinitesimal term (a value between 0 and 1, specifying the proportion of infinitesimal term relative to the total variance components including biological terms (the four categories above and infinitesimal).`
- `Noise variance component (a value between 0 and 1, specifying the proportion of noise term relative to the total variance components including all terms).`

Num_Traits [Default = 1,000]. Number of traits to be simulated.

**Population genetic parameters:**

Max_MAF [Default = 0.5]. Maximal minor allele frequency for the filtering when loading genotype file.

Min_MAF [Default = 0.05]. Minimal minor allele frequency for the filtering when loading genotype file.

Loc_Distance [Default = 0]. The minimal distance (in the unit of base-pair) between adjacent genetic variants. This is used to remove variants in strong LD with others. Since the specification of LD directly is problematic and confounded by the allele frequency, as a pragmatic solution, OmeSim uses distance to ensure sufficient independence between variants.

**Genetic model and calculation parameters:**

Iteration_Rounds [Default = 10]. Number of iterations when calculating the values of terms. Here the iterations are needed because of that the value of a term may rely on other terms whose values are not initialized at the beginning. Therefore, iteratively more and more terms will be initialized and can contribute to other terms meaningfully. The terms that only rely on genetics (and possibly infinitesimal and noise) will be finalized in the first round. The terms relying on them will be finalized in the second term. Note that, if there is a circle in the causality graph, the terms in this circle will never be finalized, however their values may be stabilized (i.e., converge to a largely unchanged value) after sufficient rounds.

Gene_Contributors [Default = 1.0, 0.5, 0.3, 0.1]. There are probabilities based on which OmeSim will randomly decide whether each gene expression contains contributions from each of the four categories of terms: `[0] = cis-genetics; [1] = trans-genetics; [2] = other gene expressions; [3] = traits.` It is suggested to specify the probability of `cis-genetics` to 1.0, mandating the presence of cis genetic contributions to the expression of the gene itself.

Traits_Contributors [Default = 1.0, 0.8, 0.5, 0.3]. There are probabilities based on which OmeSim will randomly decide whether each trait contains contributions from each of the four categories of terms: `[0] = genetics (across the whole genome); [1] = gene` `expressions; [2] = traits; [3] = infinitesimal.` It is suggested to specify the probability of `genetics` to 1.0, mandating the presence of genetic contributions to the trait.

Gene_Contri_Weights [Default = 1.0, 0.5, 4.0, 2.0]. There are the average weights to be multiplied to the values of four contributing terms (if they are present, decided by the probabilities specified in Gene_Contributors) contributing to gene expressions. The same order as in Gene_Contributors: `[0] = cis-genetics; [1] = trans-genetics; [2] = other gene expressions; [3] = traits.` Note that the weight will be automatically assigned to NaN if its term is not present.

Traits_Contri_Weights [Default = 1.0, 4.0, 2.0]. There are the average weights to be multiplied to the values of four contributing terms (if they are present, decided by the probabilities specified in Traits_Contributors) contributing to traits. The same order as in Traits_Contributors: : `[0] = genetics (across the whole genome); [1] = gene expressions; [2] = traits.` Note that the weight will be automatically assigned to NaN if its term is not present. Also note that the weight of `infinitesimal` term is not set here by weight; instead, the variance component parameter will specify the infinitesimal terms after calculating biological terms.

Weights_Relative_Range [Default = 0.5]. The relative range of the fluctuation of weights surrounding the averages specified by Traits_Contri_Weights and Gene_Contri_Weights. OmeSim will generate random weights based on the averages and the range that is allowed by this parameter.

Traits_Var_Comp_Inf_Min [Default = 0.05] Minimal variance component for the infinitesimal term (when it indeed is present). Note that this is only for traits, and gene expressions do not contain an infinitesimal term.

Traits_Var_Comp_Inf_Max [Default = 0.45] Maximal variance component for the infinitesimal term (when it indeed is present). Note that this is only for traits, and gene expressions do not contain an infinitesimal term.

Var_Comp_Noise_Min [Default = 0.1] Minimal variance component for the noise term (which is always present). Note that the non-noise part may not be deemed as "heritability" as the other traits and expressions (and noises in the traits and expressions) are also involved.

Var_Comp_Noise_Max [Default = 0.9] Maximal variance component for the noise term (which is always present). Note that the non-noise part may not be deemed as "heritability" as the other traits and expressions (and noises in the traits and expressions) are also involved.

Cis_Var_Flanking [Default = 500,00] Flanking regions (upstream or downstream) in the unit of base-pair, defining the genomic regions within which the genetic variants are considered as “cis-variants”.

Min_Var_Gene [Default = 10] The threshold of removing a gene that is too small. Here if there are fewer genetic variants in the gene (including the flanking regions specified by

Cis_Var_Flanking) than the specified value, OmeSim will remove this gene from the gene model.

Cis_Variant_Numbers_Mean [Default = 5] The average number of cis-genetic variants contributing to a gene expression.

Cis_Variant_Numbers_Range [Default = 10] The range (up or down) of the number of cis-genetic variants contributing to a gene expression. Note that OmeSim will generate the number of cis-genetic variants based on the mean and the range surrounding the mean. The minimal number of cis-genetic variant will be at least 1 (if the cis-genetic variants' contribution is indeed present, randomly decided by Gene_Contributors.)

Trans_Variant_Numbers_Mean [Default = 5] The average number of trans-genetic variants contributing to a gene expression. The same parameter also controls the genetic components for traits.

Trans_Variant_Numbers_Range [Default = 10] The range (up or down) of the number of trans-genetic variants contributing to a gene expression. Note that OmeSim will generate the number of trans-genetic variants based on the mean and the range surrounding the mean. The minimal number of trans-genetic variant will be at least 1 (if the trans-genetic variants' contribution is indeed present, randomly decided by Gene_Contributors.) The same parameter also controls the genetic components for traits.

Num_Contributing_Genes_Mean [Default = 4] The average number genes contributing to a term (either gene expression or trait).

Num_Contributing_Genes_Range [Default = 4] The range (up or down) of the number of genes contributing to a term (either gene expression or trait). Note that OmeSim will generate the number of genes based on the mean and the range surrounding the mean. The minimal number of gene will be at least 1 (if the gene expressions' contribution is indeed present, randomly decided by Gene_Contributors or Traits_Contributors).

Num_Contributing_Traits_Mean [Default = 2] The average number traits contributing to a term (either gene expression or trait).

Num_Contributing_Traits_Range [Default = 1] The range (up or down) of the number of traits contributing to a term (either gene expression or trait). Note that OmeSim will generate the number of traits based on the mean and the range surrounding the mean. The minimal number of traits will be at least 1 (if the traits' contribution is indeed present, randomly decided by Gene_Contributors or Traits_Contributors).

Num_Infinitesimal [Default = 5,000] To approximately calculate the contribution of infinitesimal term (representing the genetic background), OmeSim randomly selects a large number of genetic variants from the whole genome and add their (mostly equal and tiny) contributions. This parameter specifies how many genetic variants will be selected to form the infinitesimal term. The total value will be re-scaled based on the specified contribution of infinitesimal term.

Gene_Models [Default = 0.5,0.2,0.2,0.05,0.05] OmeSim supports five models to specify the relationship between genetic and other terms in calculating the values. They are {`additive`, `epistatic`, `compensatory`, `heterogenous`, and `compound`. Please refer to our paper for the detailed formula on how their mathematical definitions. Here, this parameter is a 5-element array to specify the probabilistic distribution of the five models among all gene expressions. `[0]=additive;[1]=epistatic;[2]=compensatory;[3]=heterogenous;[4]=compound`.

Trait_Models [Default = 0.5,0.05,0.05,0.1,0.3] The probability distribution of the above mentioned five models for traits.

Trait_Binary_Proportion [Default = 0.5] The proportion of case/control binary traits (in contrast to quantitative traits to be simulated by OmeSim. The rest (1 - Trait_Binary_Proportion) traits will be simulated as quantitative.

Binary_Mode_Proportion [Default = 0.5,0.5] Among the binary traits (proportion specified by Trait_Binary_Proportion), the proportion of traits simulated by two alternative statistical models, i.e., `liability` model and `logistic` model. The model `liability` assumes that the underlying quantitative value contributed by all terms (including biological and noise) decides the case/control status by a cutoff, above which will be diseased (i.e., case). OmeSim will identify the median of the quantitative value as the cutoff. So, half of the sample will be labeled as cases (1.0) and the other half will be controls (0.0). The `logistic` model will use the standard logistic regression model, i.e., *log(p/(1-p)) = quantitative-value*, to solve *p*, the odd (i.e., a probability) based on the quantitative values. This *p* is the probability of having the disease. OmeSim will then label the samples with odd (probability) higher than 0.5 as cases (and the rest are controls).

**Graphics genetic parameters:**

Rscript_Binary_Path [Default = "/usr/local/bin/Rscript"]. Absolute path to binary executable 'Rscript' in the executing environment. If the default doesn't work and the users is not clear on where to find the executable, please type "`> which Rscript`" to find out. Note that `R` has to be installed in the environment.

Visualization_Code_Folder. Full (absolute) path to the folder for R scripts to draw figures. The three R scripts drawing the figures are all enclosed in the OmeSim download. Note that the following libraries have to be installed before running the code (using command `install.packages('package_name')` in `R`:

- pheatmap
- tidyverse
- ggplotify
- heatmaply
- igraph

- ggraph

Large_Figures_Needed [Default = **false**]. Whether large figures for all data are needed. Generating full pictures of all data expressions and traits (correlations, causality and the original data) may be memory consuming and the outcome could be very large and difficult to view. We assume that the most useful part of the graphics function is the pathway-specific and trait-specific figures. Nevertheless, we offer this function, although the default is 'false'.

Trait_Causality_Degree [Default = 4]. Parameter to specify how many degrees between a source term (i.e., a node in the causality graph) and the target trait. The "degree" is defined as the number of edges that the source node has to go through to reach the target trait node. Sub-causality graphs for all degree < Trait_Causality_Degree will be generated.

Edge_Num_Ratio [Default = 300]. Parameter controlling the size of the sub-causality graphs. The actual size of the sub-causality graph is calculated by: Edge_Num_Ratio *Math.pow(num_of_edge_in_the_graph, 0.4).

**Parameters for the function `Causality`.**

Query_File. Full (absolute) path to a file containing lines of queries. The queries are three terms (gene names or trait names) for OmeSim to check their causal relationship via the Causality Graph. The three names should be separated using commers. Three causality relationship will be checked: "`Causality`", "`Pleiotropy`", and "`Reverse Causality`".

Figures_Needed. If elected, PNG files visualizing the causality relationship between the three terms will be generated.

Output_Folder Full (absolute) path to the folder for output files.

#### 7. Contact:

Zhou Long <>

Qingrun Zhang <>

#### 8. Copyright Licence (MIT Open Source)

Permission is hereby granted, free of charge, to any person obtaining a copy of this software and associated documentation files (the "Software"), to deal in the Software without restriction, including without limitation the rights to use, copy, modify, merge, publish, distribute, sublicense, and/or sell copies of the Software, and to permit persons to whom the Software is furnished to do so, subject to the following conditions:

The above copyright notice and this permission notice shall be included in all copies or substantial portions of the Software. THE SOFTWARE IS PROVIDED "AS IS", WITHOUT

WARRANTY OF ANY KIND, EXPRESS OR IMPLIED, INCLUDING BUT NOT LIMITED TO THE WARRANTIES OF MERCHANTABILITY, FITNESS FOR A PARTICULAR PURPOSE AND NONINFRINGEMENT. IN NO EVENT SHALL THE AUTHORS OR COPYRIGHT HOLDERS BE LIABLE FOR ANY CLAIM, DAMAGES OR OTHER LIABILITY, WHETHER IN AN ACTION OF CONTRACT, TORT OR OTHERWISE, ARISING FROM, OUT OF OR IN CONNECTION WITH THE SOFTWARE OR THE USE OR OTHER DEALINGS IN THE SOFTWARE.
